# Effective downregulation of *BCR-ABL* tumorigenicity by RNA targeted CRISPR-*Cas13a*

**DOI:** 10.1101/840900

**Authors:** Aditya Singh, Prateek Bhatia

## Abstract

**Aim:** To induce *BCR-ABL* gene silencing using CRISPR *Cas13a.*

**Background:** CML is a clonal myeloproliferative disorder of pluripotent stem cells driven by a reciprocal translocation between chromosome 9 and 22, forming a BCR-ABL fusion gene. Tyrosinekinase inhibitor drugs like imatinib are the mainstay of treatment and cases resistant to these drugs have a poor prognosis in the absence of a compatible stem-cell donor. However, with rapid advancements in gene-editing technologies, most studies are now focusing on developing a translational model targeting single-gene disorders with a prospective permanent cure.

**Objective:** To explore the potential application of the RNA targeting CRISPR-*Cas13a* system for effective knockdown of *BCR-ABL* fusion transcript in a CML cell line, K562.

**Method:** CRISPR *Cas13a* crRNA was designed specific to the chimeric BCR-ABL gene and the system was transfected as a two-plasmid system into a CML cell line, K562. The effects were enumerated by evaluating the expression levels of downstream genes dependent on the expression of the *BCR-ABL* gene. Also, next-generation sequencing was used to ascertain the effects of CRISPR on the gene.

**Results:** The CRISPR system was successfully able to lower the expression of downstream genes (*pCRKL* and *pCRK*) dependent on the activated *BCR-ABL* kinase signal by up-to 4.3 folds. The viability of the CRISPR treated cells were also significantly lowered by 373.83-fold (p-value= 0.000891196). The time-dependent kinetics also highlighted the significant in-vitro suppressive activity to last up to 8 weeks (p-value: 0.025). As per the cDNA sequencing data from Oxford MinION next-generation sequencer, the CRISPR treated cells show 62.37% suspected cleaved reads.

**Conclusion:** These preliminary results highlight an excellent potential application of RNA targeting CRISPRs in Haematological neoplasms like CML and should pave way for further research in this direction.

## 1. INTRODUCTION

Chronic myeloid leukaemia (CML) is caused by a reciprocal translocation between chromosome 9 and 22, resulting in a chimeric BCR-ABL protein having constitutive kinase activity (1). The translocation is characteristically between band q34 of chromosome 9 bearing Abelson (ABL) proto-oncogene and band q11 of chromosome 22 bearing the breakpoint cluster region (BCR) gene (2). The BCR-ABL fusion gene protein can vary from 190 kDa to 230 kDa depending upon the breakpoint site on BCR gene (3). Most commonly found BCR-ABL fusions carry their breakpoints in the 2.9 kb region between exons 13 and 15 (b2 and b4) of BCR region that fuses with the large intronic region of ABL between exons 1b and 2b. This forms an 8.5 kb BCR-ABL transcript containing BCR exon b2/b3 and ABL exon 2 (a2). Both combinations (b1a2 and b3a2) code for the same 210 kDa protein (3). However, irrespective of the breakpoint, the BCR-ABL chimeric kinase phosphorylates and activates downstream proteins like pCRKL, pCRK, pSTAT5 etc, that further modulate expression of target genes to give proliferative advantage to the Philadelphia chromosome (Ph) carrying pluripotent stem cells (4).

With recent advancements in gene editing research, much focus has drifted towards targeting single gene disorders for possible therapeutic cure. In this regard, Clustered Regularly Interspaced Short Palindromic Repeats (CRISPR), has emerged as a highly versatile, flexible and cost-effective tool for gene editing experiments. This tool has circumvented the shortcomings of the previous tools like transcription activator-like effector nucleases (TALENs) and zinc-finger nucleases (ZFNs) with improved overall turnaround time (5–7). In haematological neoplasms, the chimeric fusion kinase BCR-ABL of CML seems an exciting target for CRISPR as it is the driving force behind the neoplastic pathogenesis. Working on the same idea, in 2017, García-Tuñón et al, demonstrated the use of CRISPR Cas9 for targeting BCR-ABL chimeric trans-gene (8) in a synthetically engineered Boff-p210 cell line bearing the transgene coding for the p210 variant of the BCR-ABL fusion gene. This extensive work showcased for the first time the application of CRISPR in downregulating the BCR-ABL chimeric gene. However, in actual cases of CML, the breakpoints might differ at the DNA level due to different intronic breakpoint regions in ABL gene (9). Hence, using DNA targeting CRISPR will require a per case targeted sequencing and custom CRISPR sgRNA/ sgRNAs to effectively knock down the BCR-ABL chimeric gene. This would require a dedicated research team which will eventually incur substantial costs and increase the turnaround time per case. Also, the custom target effectiveness might vary over the range of patients and hence a definitive prognosis might never be established. All these issues can be circumvented by employing techniques that can target the BCR-ABL transcript rather than the DNA as the former will remain immune to the intronic changes at the DNA level. This was only possible after the recent development and assessment of RNA targeting CRISPRs (10). These CRISPR systems, unlike DNA targeting Cas9, are additionally more flexible and exhibit less off-target and more on-target activities. In the current experimental study, we’ve explored the use of CRISPR Cas13a, a form of RNA targeting CRISPR, in knocking down the BCR-ABL transcript and quantified the downstream gene products (pCrKL and pCrK) using flow-cytometry and western blot techniques, resulting in a significantly large reduction in the viability of the treated K562 cells. The overall activity of CRISPR has been enumerated by deep sequencing of cDNA by Oxford Nanopore Technologies MinION system. The developed CRISPR system was found to be effectively downregulating the levels of downstream genes of the CML pathogenesis pathway. Moreover, we have tried highlighting the time-dependent kinetics of the downstream effects of the editing. To the best of our knowledge, this is the first such in vitro experimental study successfully employing RNA targeting CRISPR-Cas13a in knocking down the BCR-ABL transcript and is likely to pave way for further research in the field.

## 2. MATERIALS AND METHOD (FOR RESEARCH ARTICLES ONLY)

The methodology is briefly illustrated in form of a flowchart for in figure 1.

**Figure 1:**
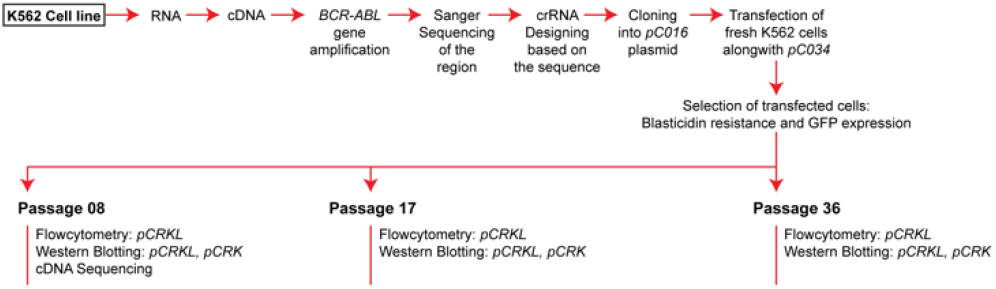
Flowchart of the study design

### 2.1 Construction CRISPR plasmids

The CRISPR RNA (crRNA) was designed using CRISPR-RT server (11). Only the crRNA designs overlapping *BCR* and *ABL* breakpoint region without any predicted off-targets were considered. The top and bottom crRNA oligos with *BbsI* overhangs were procured from Sigma-Aldrich (St. Louis, Missouri, US). The two oligos were phosphorylated using T4 polynucleotide kinase and ligated using T7 polymerase to form a double-stranded crRNA as per the protocol described by Cong *et al,* 2014 (12). The plasmids *pC016 - LwCas13a guide expression backbone with U6 promoter* (Addgene #91906) and *pC034 - LwCas13a-msfGFP-2A-Blast* (Addgene #91924) were obtained as gifts from Dr Feng Zhang. The plasmid pC034 carries blasticidin resistance and green fluorescent protein (GFP) expression markers. The plasmid pC016 was digested using *BbsI-HF* (New England Biolabs, MA, USA, Cat#R3539L) as per the manufacturer’s protocol with a slight modification of over-digesting the plasmid in 10U of the enzyme thrice in continuity with 16hrs of incubation each time without deactivating the enzyme in between ensuring a successful second cut after the plasmid is linearized by the first cut. The pC016 plasmid with insert (pC016I) was then transformed into competent *Escherichia coli DH5a* cells using the heat shock method as described by Sambrook and Russell (13). The plasmid was extracted from the positive clone using QIAprep Spin Miniprep kit as per the manufacturer’s protocol.

### 2.2 Transfection of K562 cells with CRISPR plasmids

The K562 cell line used in the study is a Ph chromosome-positive CML cell line which was first described in 1975 by Lozzio CB and Lozzio, BB (14). The characteristic *BCR-ABL* sequence was sequenced and verified before proceeding with the experiments. Lipofectamine LTX (cat#15338100, Thermo-Fisher Scientific, MA, US) was used for the co-transfection of the two plasmids, pC016I and pC034 using the manufacturer’s protocol in a six-well plate containing approximately 50,000.00 cells from an overnight culture of K562 cells in RPMI-1640 medium with 1% Penicillin-Streptomycin at a confluency of 75-80%. The cells were incubated overnight at 37°C with 5% CO2. After the overnight incubation, old media was replaced with fresh and additionally 6 μg/mL of blasticidin antibiotic was added as the positive selection marker for the plasmid in both test and negative control cells (15). Cultures were observed until all negative control cells were killed and GFP expression of the transfected cells became close to a hundred per cent. Cells were harvested at passage number eight, seventeen and thirty-six for downstream analysis and time-dependent kinetics study.

### 2.3 Downstream gene regulation analysis

Total RNA was extracted using the RNeasy kit from QIAGEN (cat#74104) as per the manufacturer’s protocol. A fraction of RNA was reverse transcribed to cDNA using RevertAid first-strand cDNA synthesis kit by Thermo Fisher scientific (cat#K1622) using the manufacturer’s protocol. The *BCR-ABL* cDNA region was amplified using specific primers, forward 5’-ACAGAATTCCGCTGACCATCAATAAG-3’, reverse 5’-TGTTGACTGGCGTGATGTAGTTGCTTGG-3’ (16). Selection of target downstream genes was done after consulting with the CML pathway from KEGG with pathway ID hsa05220. We found that the genes *CRK* and *CRKL*, among a few others, are directly phosphorylated by the expression of BCR-ABL chimeric gene. We chose these two genes, *pCRK* and *pCRKL* denoting the phosphorylated forms of them for the downstream expression analysis. For the flow cytometric analysis of *pCRKL* levels, the *pCRKL (Y207)* specific Alexa Fluor 647 tagged antibody (cat#560790, Becton, Dickinson and Company, NJ, US) was used as per the manufacturer’s protocol in a Beckman Coulter Gallios flow cytometer (Beckman Coulter Inc., IN, US). The fold change was calculated by comparing the mean fluorescence intensity (MFI) using median population and at least 10,000 events per test between K562 cells as control and K562 cells bearing CRISPR plasmids (coded as K562-1634). The calculations were done in Kaluza Analysis software by Beckman Coulter.

Whole-cell protein extraction for western blot was done using Radioimmunoprecipitation assay (RIPA) buffer (cat#89900, Thermo Fisher Scientific, MA, US) using the manufacturer’s protocol. Bicinchoninic acid (BCA) protein assay was done for extracted protein quantification using Pierce BCA Protein assay kit according to the manufacturer’s protocol (cat#23225, Thermo Fisher Scientific, MA, US). For western blotting, primary rabbit raised antibodies against *pCRKL Y207* (37 kDa) and *pCRK Y221* (34 kDa) (another downstream target protein) were procured from Abcam plc, Cambridge, the UK with catalogue numbers ab52908 and ab76227 respectively and housekeeping mouse raised *β-tubulin* (51 kDa) primary antibody was procured from Cell Signalling Technologies, MA, the US with catalogue number 2146S. The anti-rabbit and anti-mouse secondary horseradish peroxidase linked antibodies were procured from Cell Signalling Technologies with catalogue numbers 7074P2 and 7076P2 respectively. Image J software was used for the analysis of the western blot imaging data (17).

### 2.4 Cell viability assay

The cell viability assay was performed in triplicate using Promega *CellTitre-Glo* 2.0 reagent as per the manufacturer’s protocol for establishing the role of knockdown of *BCR-ABL* transcript on the cells.

### 2.5 Enumeration of CRISPR activity by NGS

Total RNA from K562-1634 and K562 cells were subjected to cDNA sequencing using Oxford Nanopore Technologies MinION instrument with SQK-PCS108 library kit along with barcodes from SQK-PBK004 kit on an FLO-MIN106 R9.4 flow cell according to the manufacturer’s protocol. QCAT tool was used for demultiplexing the barcoded reads (https://github.com/nanoporetech/qcat), NanoFilt was used to filter the data with a minimum quality cut off of 5 (18). NCBI local BLAST+ was used to extract raw reads carrying crRNA sequence (19). The reads carrying the crRNA sequence were extracted from the whole reads using NCBI Basic alignment search tool (BLAST) (19) and then aligned with *BCR-ABL* cDNA and crRNA sequences using BWA-MEM(20). The resulting SAM alignment format was coordinate sorted and indexed using SAMtools (21) and visualized in Integrative Genomics Viewer (IGV) (22). The study was duly approved by the Institute’s Ethics Committee and the Departmental Review Board, Postgraduate Institute of Medical Education and Research, Chandigarh, India vide the letter-number INT/IEC/2017/44.

### 2.6 Statistical analysis

SPSS Statistics software version 25 (IBM Corp, NY, US) was used to perform statistical tests. The flow cytometry and western blot data were normalized using maximum value before performing any statistical tests. To achieve this, the value of the sample having the maximum reading for the experiment was used to divide the values of all the samples from that experiment. This was done to bring all the values between zero to one, to enable us in making a common trend graph for expression of the genes independent of the method used (flow cytometry or western blot). Student’s t-test was used to check for significant change in expression over the period. Pearson’s correlation was used to find a significant relation between flow and western blot data.

## 3. RESULTS (FOR RESEARCH ARTICLES ONLY)

### 3.1 crRNA Designing

CRISPR-RT suggested a total of 175 target regions, out of which, 14 were overlapping the *BCR-ABL* chimeric region. Among these 21 candidate crRNAs, the one 5’-ATTTAAGCAGAGTTCAAAAGCCCTTCAG-3’, which spanned significantly in both *BCR* and *ABL1* regions (17 and 12 bases in *BCR* and *ABL1* respectively) with the lowest GC content of 39% and no predicted off-targets was chosen (Figure S1). There were no predicted off-targets because *Cas13a* tolerates only one mismatch and in case of our crRNA design, the resulting CRISPR won’t be able to target wild type BCR or ABL because of 12 and 17 bases mismatch respectively.

### 3.2 Transfection and downstream analysis

Pure colonies of K562-1634 transfected cells were selected by day 12 (Figure 2). Flow cytometric analysis was done in duplicate for each run and the average was used for calculations. At least ten thousand events per reaction were recorded with maximum events recorded reaching around 85,000 in a single reaction. *pCRKL* level was observed to be 4.30, 4.02 and 1.80 times lower than the control K562 cell levels at passages eight, seventeen and thirty-six respectively (p-value 0.025) (Figure 3A). Western blotting data of *pCRKL* and *pCRK* (Figure 2 B and unedited pictures in figures S2-S4) was found to be synchronous with the flow cytometry data after normalization with a significant p-value of 0.01 (Figure 3 C). Cell viability assay performed for passage 11 cells in triplicate for control and treated cells after compensating the blank readings were 1300479, 1237852, 1173013 and 3,378, 2974, 3,576 respectively. This translates into a 373.83-fold (p-value: 0.000891196) reduction in the viability of the cells when treated with CRISPR (Figure 4).

**Figure 2:**
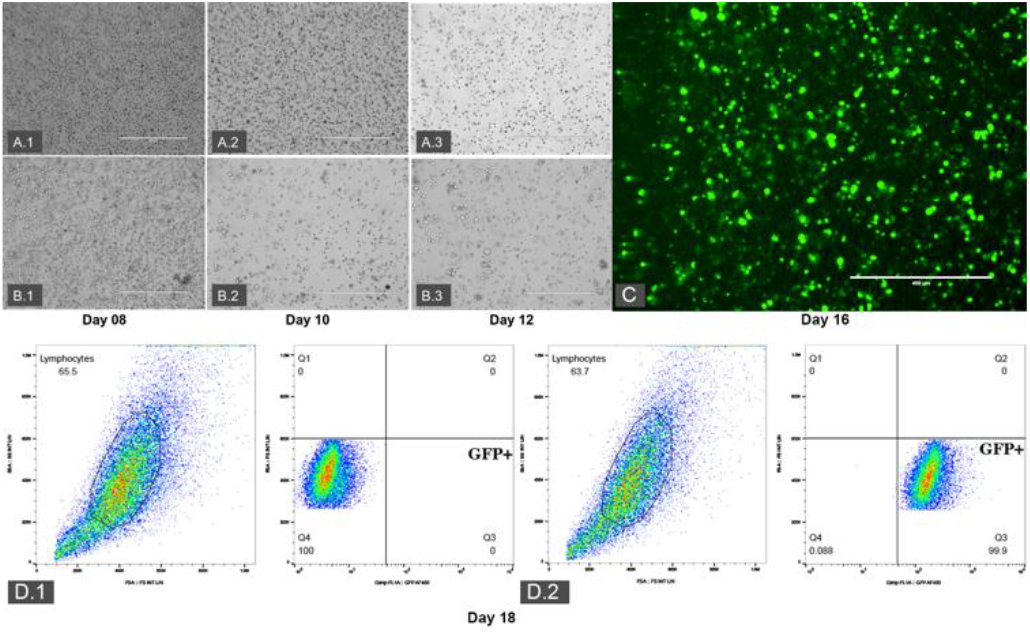
Blasticidin selection of K562-1634 cells over different passages in contrast with wild type K562 cells. A1-A3 and B1-B3 Inverted microscopy images taken at 10x magnification on days 08, 10 and 12 for control K562 and 1634 plasmid bearing K562 cells respectively. C: Final resulting cell population of 1634 plasmid bearing cells at 10x magnification showing widespread GFP expression. D: Flowcytometric estimation of GFP positive cells in control (D.1) cells not subjected to blasticidin and transfected (D.2) cells at day 18.

**Figure 3:**
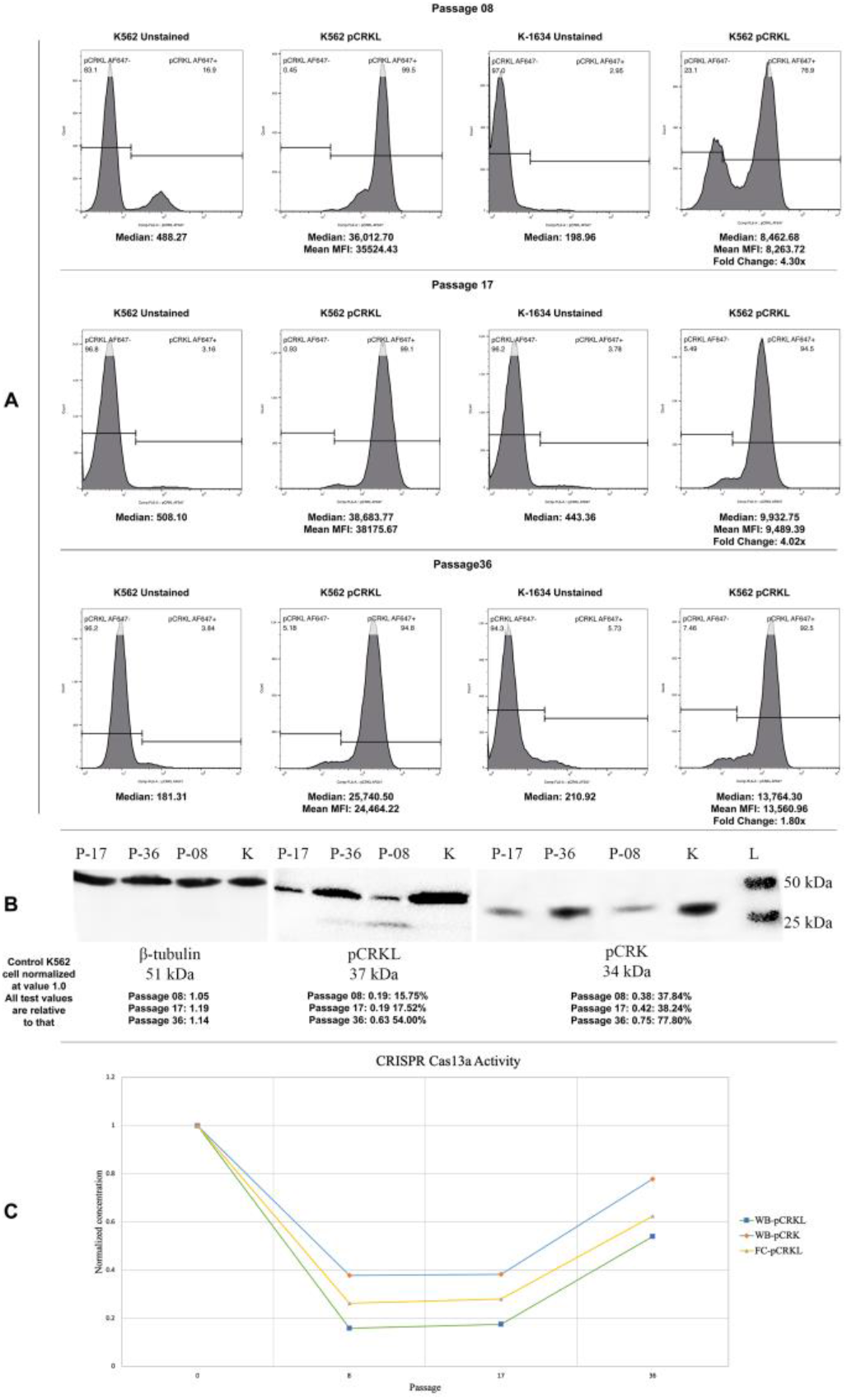
Flow cytometry and western blot data for control vs K562-1634 cells. A: Flow cytometry data for comparative pCRKL expression levels. K-1634 denotes the K562 cells bearing the CRISPR plasmids. B: Comparative western blot data for pCRK and pCRKL expression levels. K-K562 cells without plasmids (control). P08, P17 & P36 denote passage numbers 8, 17 and 36 for K562 cells bearing CRISPR plasmids (K-1634 cells). C: Comparative trends between western blot and flow cytometry data after normalization.

**Figure 4:**
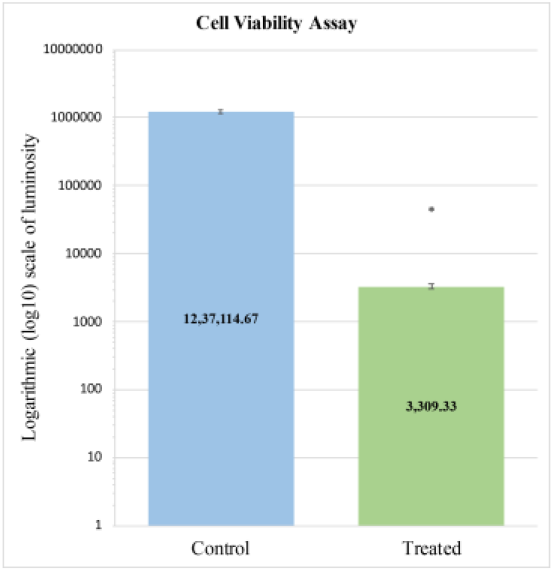
Cell viability comparative assay between untransfected K562 control and CRISPR treated K562 cells (K-1634).

### 3.3 NGS Sequencing of transfected and control cell lines

The next-generation sequencing using Oxford MinION generated a total of 5,365,731 reads with an average read length of 402.8 bp, median read length of 253 bp and maximum read length of 232,778 bp. As far as the calculation of the CRISPR activity, we found all the reads to be shorter than the total reads on in the alignment data but since the NGS reads are short and incomplete, only the reads which were found to begin and end abruptly around 112 bp in the 257 bp focus region of the *BCR-ABL* transcript upstream of the crRNA region were accounted for CRISPR activity. According to this, out of the 93 reads, 58 reads were cleaved by the CRISPR activity, which translates to a 62.37% activity (Figure S5). Interesting to note here was the fact that the reads also show base substitutions and INDELs in the crRNA region but *Cas13a* is not known to cause them hence we’ve associated them to the platform errors of the sequencer.

## 4. DISCUSSION

There are four major genome editing tools, namely, Meganucleases (MNs), ZFNs, TALENs and the most recent, CRISPR. CRISPR finds its edge in being almost 10 times cheaper than its closest competitor, TALENs, has widest applications, is freely available and simplest to design and apply (23). Earlier, in a study by García-Tuñón *et al,* it was observed that CRISPR *Cas9* was able to effectively control the oncogenic capabilities of *BCR-ABL* chimeric gene in custom Boff-p210 cell line carrying *BCR-ABL* fusion transgene (8). In a recent study, Liu *et al* demonstrated effective downregulation of *BCR-ABL* gene using CRISPR RNA guided *Fokl* Nucleases in both cell lines and murine xenograft model (24). However, García-Tuñón *et al,* in their other research article on CRISPR, published recently, highlighted the limitation of CRISPR/Cas9 in effectively knocking out target DNA without a null variation inducing homology-directed repair template in the construct. They went on to demonstrate that effective knockdown could be achieved if sgRNAs target the splice donor site of the gene (25). Moreover, it has been shown that in CML, using a DNA targeting CRISPR would mean designing custom guide RNAs per patient as the breakpoints might differ per case at the DNA level (9).

Considering above limitations and problems in targeting *BCR-ABL* gene at the DNA level by CRISPR/Cas9, we hypothesized utility of targeting the p210 transcript of CML with an RNA CRISPR since the p210 transcript wouldn’t be likely affected by the differences in intronic breakpoints. Working on this hypothesis, we designed this study to explore the potential of RNA targeting CRISPR *Cas13a* in knocking down the tumorigenicity of the *BCR-ABL* chimeric gene. CRISPR *Cas13a* is a relatively newer form of CRISPR which targets RNA rather than the DNA and has been shown to have an efficiency of around 50% (10). In the current study, we found the efficiency to be around 62.37%. This RNA targeting CRISPR system was able to lower the expression of downstream genes of CML pathogenesis as per the KEGG (26) pathway ID hsa05220.

We used *pCRKL* and *pCRK* as surrogate markers for enumerating the knock-down of *BCR-ABL* as their expression is directly proportional and dependent on the expression of it. *pCRKL* levels as calculated through passage numbers 8, 17 and 36 were 23.26, 24.85 and 55.43% by flow cytometric analysis and 15.75, 17.52 and 54.00% respectively by western blotting experiment. The same for *pCRK* was found to be 37.84, 38.23 and 77.80% through passage numbers 8, 17 and 36 respectively as per the western blotting data. This suggests that the activity of CRISPR was only sustained up till passage number 17 and beyond that the cells had lost the plasmid which was evident after incubating the cells with blasticidin and observing that most of the cells were killed in few days.

The cell viability assay confirmed that the cells were significantly less viable than the control cells. This was because the *BCR-ABL* gene provides a proliferation advantage to the cells and when its transcripts were knocked down by CRISPR, it also led to the drop in the viability of the cells. On the analysis of the *BCR-ABL* transcript changes by NGS, we could observe possibly truncated reads upstream of the crRNA region, which is a known mechanism as described by Wolter, F. et al., in 2018 (27). Overall, our study results show that CRISPR-*Cas13a* efficiency (based on NGS) and downregulation of downstream genes (*pCRKL* & *pCRK*) go hand in hand *in vitro*.

It is interesting to mention here that imatinib in a dose-dependent manner of 10 ≥μM reduces *pCRKL* levels to 16.6% which translates to a 6.05-fold reduction (28), compared to the 4.3-fold reduction achieved by the described CRISPR *Cas13a* RNA machinery. However, as detailed above, with rapid advancements in the CRISPR technology, this gap might be filled or even surpassed in the near future. The next phase for the system once fully developed and established is the delivery into actual patients. As *Cas* is a bacterial protein, it does pose some hurdles, mainly immunological once we try to deliver the same into actual patients (*in vivo*). There are delivery methods such as viral vectors, ribonuclear complexes or plasmid DNA but these all have certain drawbacks like a viral vector might integrate the *Cas* gene into the host’s genome making it produce the protein continuously, hence, giving ample time for off-target effects to manifest. Also, permanently changing the host DNA is a risk which we do not currently have complete knowledge to fathom. There are also elevated immune responses in the case of viral vectors. Currently, the best possible way to deliver these components is in form of synthetic ribonuclear protein (RNP) complexes which are transient, do not cause much of immune response and is easier to upscale in pharmaceutically. These RNP complexes are composed of both the crRNA and Cas protein (29).

## CONCLUSION

The study is limited by further analysis on primary culture of multiple CML patients. Utilizing newer, more efficient RNA targeting CRISPRs like CRISPR *Cas13b* and *Cas13c* with efficiencies going up to *90-95%* (30) can potentially revolutionize the effects further with higher fold reduction in downstream target protein expression, ultimately reducing the tumorigenic capabilities of *BCR-ABL* chimeric gene. The described work acts as a proof of concept and paves way for CRISPR-based studies in hematological neoplasms like CML which are an excellent target for it.

## Supporting information

Graphical abstract

## ETHICS APPROVAL AND CONSENT TO PARTICIPATE

Not applicable.

## HUMAN AND ANIMAL RIGHTS

No animals were used as the basis of this study. This research was conducted according to the Declaration of Helsinki principles.

## CONSENT FOR PUBLICATION

Not applicable.

## AVAILABILITY OF DATA AND MATERIALS

Not applicable.

## ACKNOWLEDGEMENTS

AS would like to thank Dr Meenu Singh for her valuable insights during the work.

## Supportive/Supplementary Material

**Figure S1:** Alignment of crRNA candidate sequences with BCR-ABL sequence.

**Figure S2:** Unedited raw western blot image for *β-tubulin* estimation

**Figure S3:** Unedited raw western blot image for *pCRKL* estimation

**Figure S4:** Unedited raw western blot image for *pCRK* estimation

**Figure S5:** cDNA reads aligned to reference *BCR-ABL* region from the Oxford MinION sequencing data. A: Control K562 reads. B: Reads from K562 cells bearing the CRISPR plasmids showing cleaved reads.

