## Supplementary figures and images for "Effective downregulation of *BCR-ABL* tumorigenicity by RNA targeted CRISPR-*Cas13a*"

### Figure S1

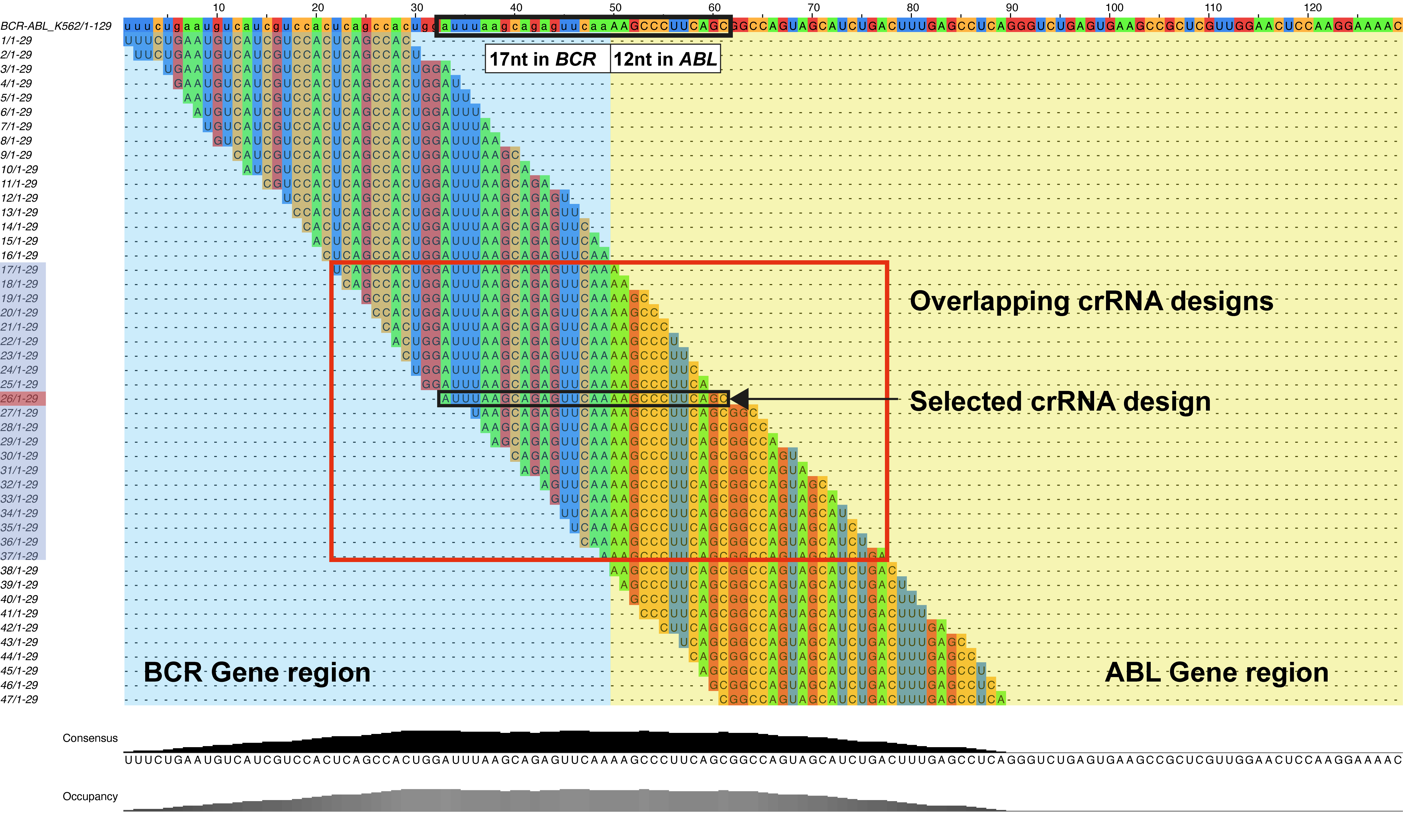

### Figure S2

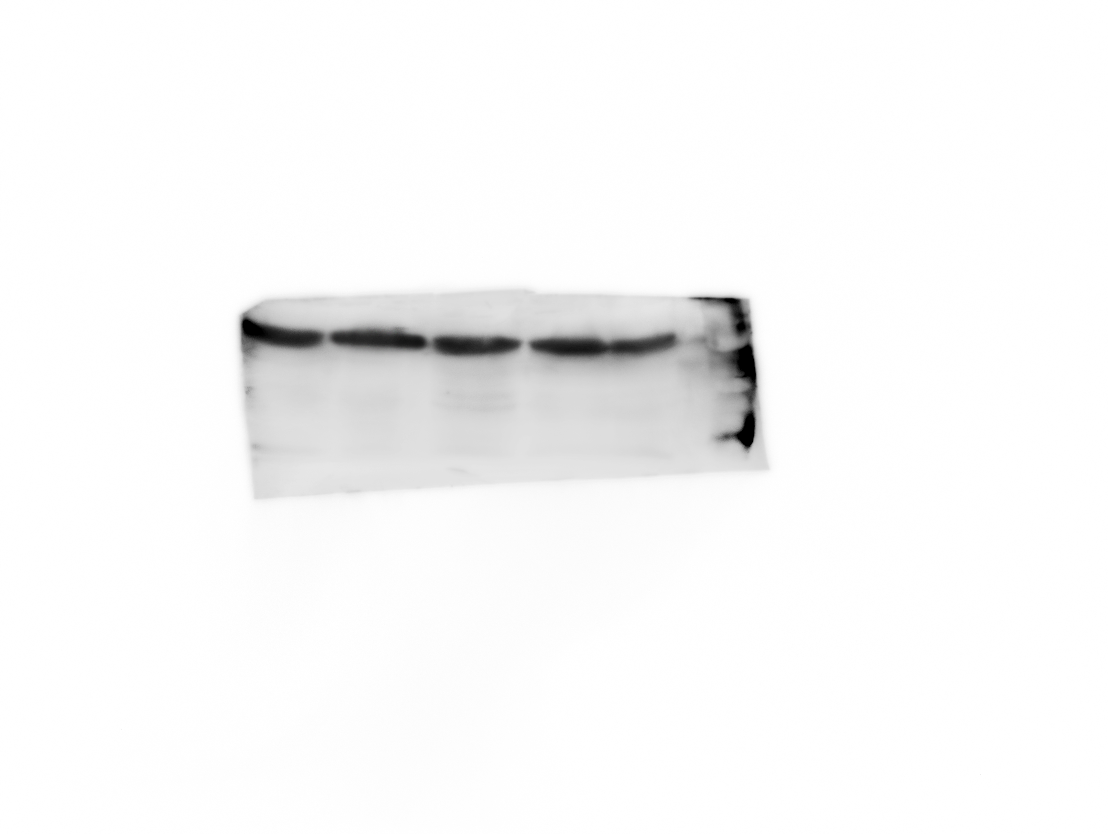

### Figure S3

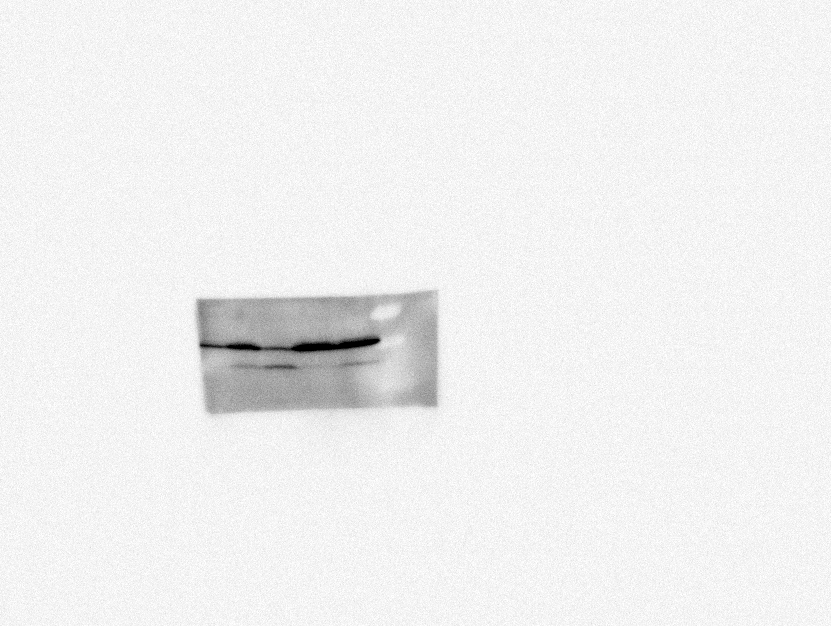

### Figure S4

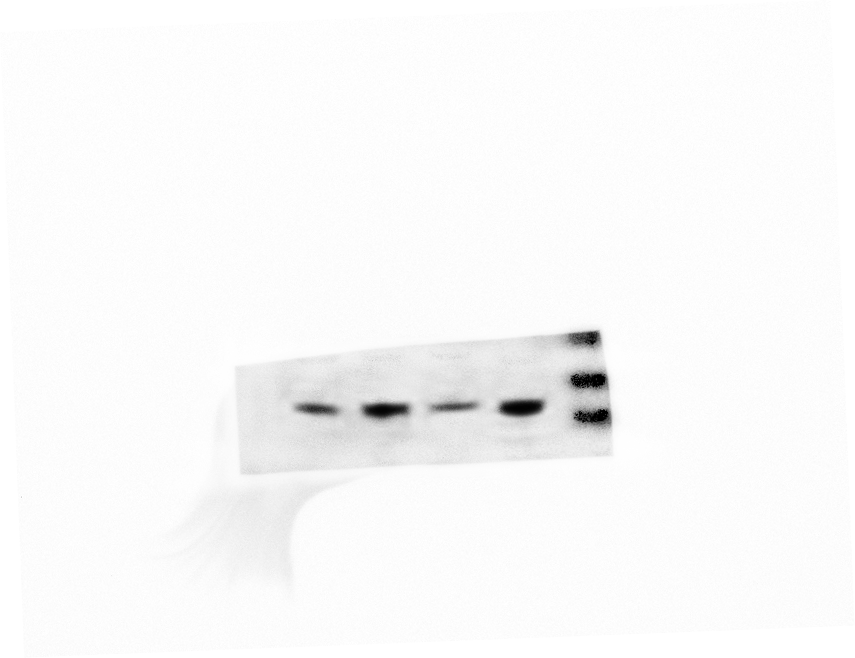

### Figure S5

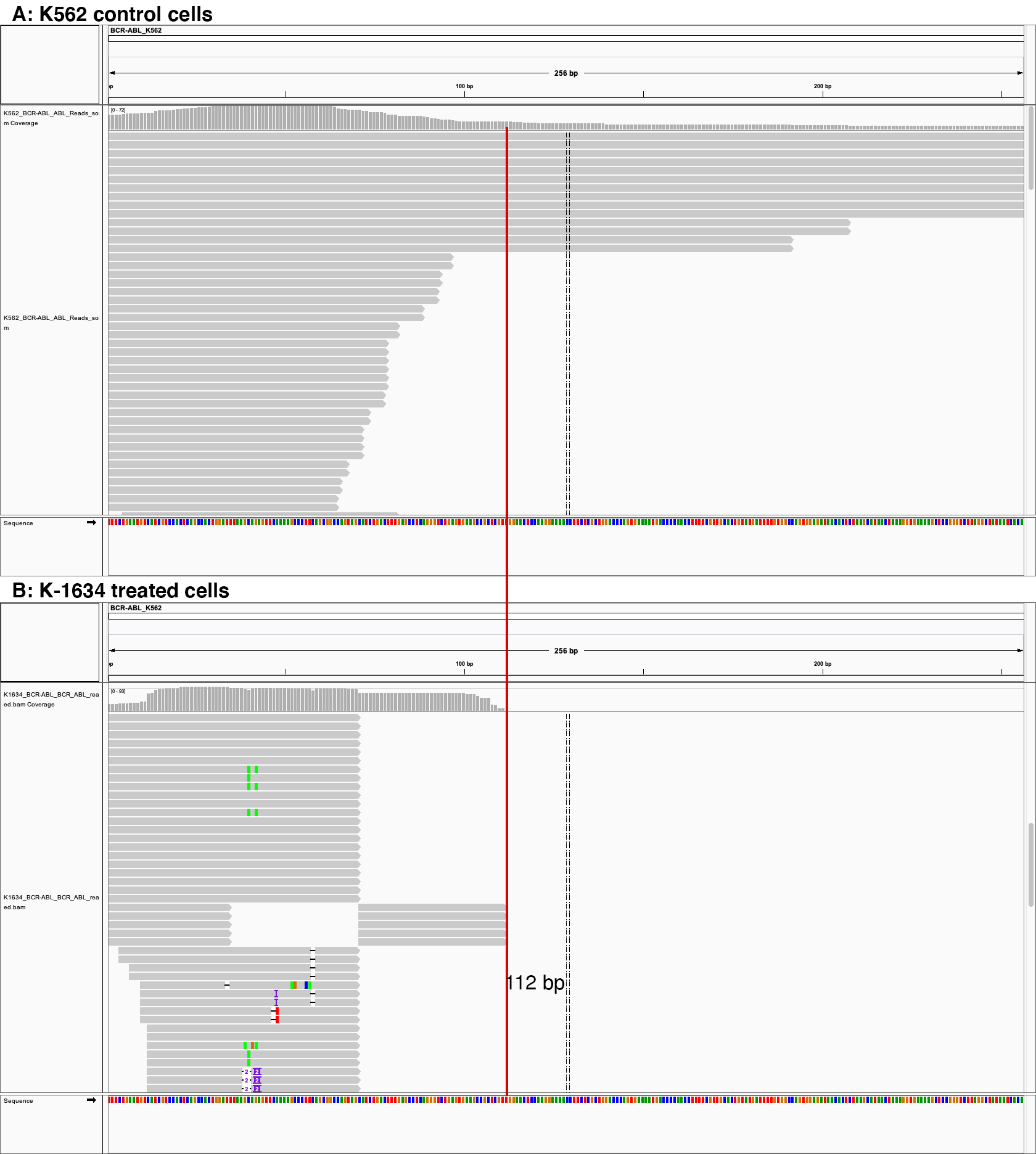

### Graphical abstract

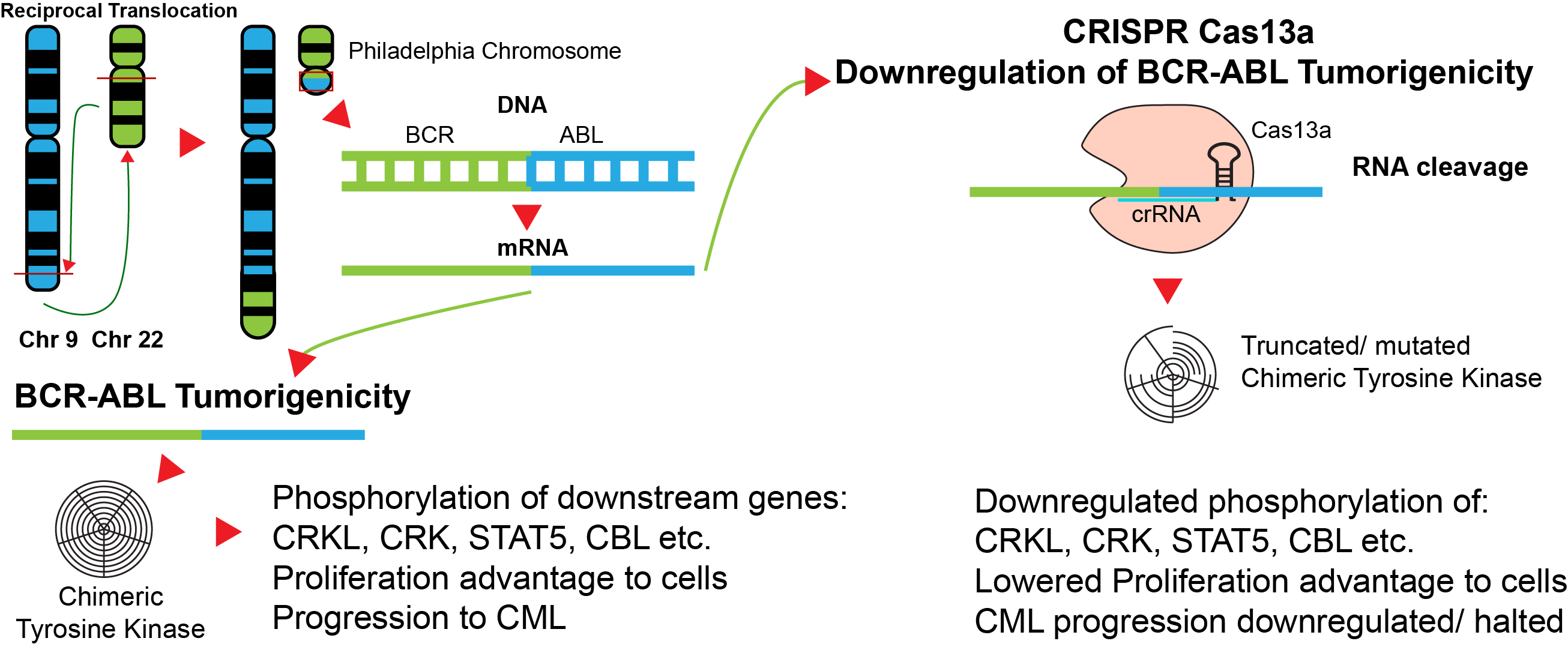
